# Applied tutorial for the design and fabrication of biomicrofluidic devices by resin 3D printing

**DOI:** 10.1101/2021.11.23.468853

**Authors:** Hannah B. Musgrove, Megan A. Catterton, Rebecca R. Pompano

## Abstract

Stereolithographic (SL) 3D printing, especially digital light processing (DLP) printing, is a promising rapid fabrication method for bio-microfluidic applications such as clinical tests, lab-on-a-chip devices, and sensor integrated devices. The benefits of 3D printing lead many to believe this fabrication method will accelerate the use of microfluidics, but there are a number of potential obstacles to overcome for bioanalytical labs to fully utilize this technology. For commercially available printing materials, this includes challenges in producing prints with the print resolution and mechanical stability required for a particular design, along with cytotoxic components within many SL resins and low optical compatibility for imaging experiments. Potential solutions to these problems are scattered throughout the literature and rarely available in head-to-head comparisons. Therefore, we present here a concise guide to the principles of resin 3D printing most relevant for fabrication of bioanalytical microfluidic devices. Intended to quickly orient labs that are new to 3D printing, the tutorial includes the results of selected systematic tests to inform resin selection, strategies for design optimization, and improvement of biocompatibility of resin 3D printed bio-microfluidic devices.

## 1. Introduction

In microfluidic device development, a recurring theme is to complete bioanalytical assays at a fraction of the time and cost required for macroscale methods. This aspiration makes rapid and accessible fabrication of microfluidic devices a key goal. Historically, microfluidic fabrication relied heavily on soft lithography methods such as casting polydimethylsiloxane (PDMS) on hard micropatterned masters.^1,2^ Soft lithography is primarily a 2D fabrication method, in which multi-layer devices are generated by tedious or challenging manual alignment and bonding. This process is sensitive to any dust that falls onto the layers during aligning and bonding, especially if conducted outside a cleanroom environment. Thus, in order to produce complex 3-dimentional devices monolithically and without a clean room, many groups have turned to 3D printing as a simplified workflow for rapid fabrication in the laboratory.^3–6^

Since the early 2010s, stereolithographic (SL) 3D printing has emerged as a promising technique for fabricating microfluidic devices.^5,7,8^ Briefly, this technique works by curing photopolymerizable resins with a UV or visible light source in sequential layers that build on top of one another. Stereolithographic apparatus (SLA) printers were some of the first SL printers and utilize a UV laser guided by mirrors, curing resin point-by-point in a scanning manner in the x and y directions. Direct light processing (DLP) printers were developed later and utilize UV projectors that allow an entire layer to be cured simultaneously from a direct light path. Because of their direct light path, DLP printers tend to have slightly better resolution compared to the similar SLA printers.^5,9^ The development of higher resolution DLP printing, along with other SL printers, has increased the use of 3D printing in fields of dentistry, audiology, medicine, and microfabrication.

Compared to traditional fabrication methods, 3D printing reduces cost and fabrication time while increasing product customization.^5,10^ Having the ability to readily customize microfluidics also expediates the “fabrication-application-results” process for devices.^1,11^ Indeed, many bioanalytical 3D printed devices have been well documented in reviews.^12–16^ Applications have ranged widely from bioreactors and probes made for direct contact with a range of cell lines,^17–20^ to approaches that utilized 3D-printed molds to cast microfluidic devices in other materials.^21–23^

Like all fabrication methods, 3D printing requires compromises between desirable features, e.g. resolution and cytocompatibility, and the past 5 years have seen an explosion of papers seeking to address key limitations. The ability to print leak-free devices, small internal microchannels (<1000 μm), and biocompatible, optically-clear devices depends on several factors that will be discussed further below, including aspects that affect mechanical integrity and the rate and resolution of photopolymerization (Fig.1).^24–26^ For commercial resins, strategies to diminish resin cytotoxicity have started to emerge recently,^27–30^ along with solutions addressing print resolution,^31–36^ imaging compatibility,^33,37–39^ and surface modification and bonding.^40,41^ Sifting through the plethora of literature for best practices can be challenging for researchers who are new to this rapidly growing field.

**Fig. 1.**
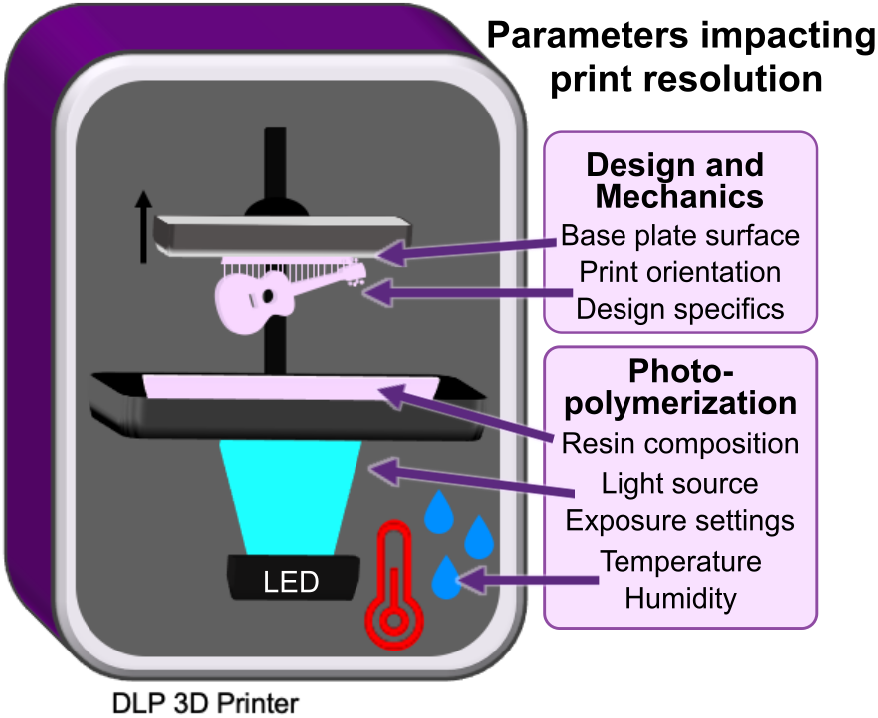
Schematic of a DLP 3D printer, highlighting the design and mechanical factors as well photopolymerization parameters which influence print resolution. Digital light processing 3D printers project UV or violet light through optically clear sheets (usually Teflon) into a vat of photopolymerizable resin (pink). In locations where the light is projected, the resin crosslinks to form a solid structure. Exposure and crosslinking are performed layer by layer on the base plate, which lifts up as each concurrent layer is formed. Production of a clean print is dependent on instrumental, environmental, chemical, and design elements that impact either the print surface (base plate), mechanics (print orientation, design specifics), or chemical reaction (resin composition, light source, exposure settings, temperature, or humidity).

Here we present an applied, data-supported guide to the key factors that should be considered for design and fabrication of SL 3D printed biomicrofluidic devices, written especially for groups that are new to this growing field. Types of instrumentation and resin formulations have been reviewed and characterized in depth recently, and will not be presented in detail here.^8,16,42–44^ This tutorial follows the order of a typical workflow by first considering resin selection and then demonstrating how to increase print resolution by using simple changes in feature design and printer settings. Following printing, cytotoxicity of materials is addressed, particularly the extent and effectiveness of post-treatment strategies for applications involving contact with primary cells. Finally, preparation and considerations for use with fluorescent microscopy is outlined with data displaying autofluorescence and optical clarity of materials. We include the results of systematic, head-to-head comparisons of printing and post-treatment conditions designed to optimize the integrity of a printed piece, the resolution of interior channels, and the biocompatibility of the part. It is our hope that this work will streamline the adoption of 3D printed devices by more specialized biomedical research fields, bioanalytical laboratories, and others new to 3D printing.

## 2. Materials and Methods

### 2.1 3D Design and Printing

All printed pieces used for this work were designed either in Autodesk Fusion 360 2020 or Autodesk Inventor 2018 and exported as an .stl file. DWG files of prints shown in figures can be found in the supplementary information. Files were opened in the MiiCraft Utility Version 6.3.0t3 software, where pieces would be converted into sliced files with appropriate layer heights. The files were converted to the correct file format to include settings optimized for either the CADworks3D M50−405 printer (MiiCraft, CADworks3D, Canada), which had a 405 nm light source, or for the CadWorks 3D Printer P110Y, which had a 385 nm source. All prints were printed in one of three resins: FormLabs BioMed Clear V1 (FormLabs, USA), FormLabs Clear V4 resin (FormLabs, USA) or MiiCraft BV007a Clear resin (CADworks3D, Canada). These resins were chosen as representative of those formulated for biocompatibility (FL BioMed), standard clear printing (FL Clear), or high-resolution microfluidics (MC BV007a).

After printing, all materials were rinsed with isopropyl alcohol using the FormWash from FormLabs, following the wash suggestions from the FormWash online guide.^45^ Alternatively, materials were placed in a container with IPA and placed on a rocker for longer periods, extending times by 5 minutes for low viscosity materials and by 1 hour for more viscous resins. Once residual resin was rinsed from the prints, the pieces were dried with compressed air and post-cured with additional UV dosages using the FormCure (FormLabs, USA). Specific print and post-print settings for each resin and printer can be found in Table S1.

### 2.2 Viability testing with primary murine lymphocytes

#### 2.2.1 Primary cell preparation

Animal work was approved by the Institutional Animal Care and Use Committee at the University of Virginia under protocol #4042 and was conducted in compliance with guidelines from the University of Virginia Animal Care and Use Committee and the Office of Laboratory Animal Welfare at the National Institutes of Health (United States). Following isoflurane anesthesia and cervical dislocation, spleens were harvested from female and male C57BL/6 mice between 8-12 weeks old. The spleens were collected into complete media consisting of RPMI (Lonza, Walkersville, MD, USA) supplemented with 10% FBS (VWR, Seradigm USDA approved, Radnor, PA, USA), 1× l-glutamine (Gibco Life Technologies, Gaithersburg, MD, USA), 50 U/mL Pen/Strep (Gibco, MD, USA), 50 μM beta-mercaptoethanol (Gibco, MD, USA), 1 mM sodium pyruvate (Hyclone, Logan, UT, USA), 1× non-essential amino acids (Hyclone, UT, USA), and 20 mM HEPES (VWR, PA, USA).

To produce a splenocyte suspension, harvested spleens were crushed through a 70-μm Nylon mesh filter (Thermo Fisher, Pittsburgh, PA, USA) into 10 mL of complete media. The cells were then centrifuged for 5 minutes at 400 xg. The pellet was resuspended into 2 mL of ACK lysis buffer, which consisted of 4.15 g NH_4_CL (Sigma-Aldrich, St. Louis, MO, USA), 0.5 g KHCO_4_ (Sigma, MO, USA), 18.7 g Na_2_EDTA (Sigma, MO, USA) into 0.5 L MiliQ H_2_O (Millipore Sigma, Burlington, MA, USA). The cells were lysed for 1 minute before being quenched by bringing up the solution to 10 mL with complete media and immediately centrifuging again. The pellet was resuspended into 10 mL of complete media, and density determined by trypan blue exclusion. To prepare for cell culture, the suspensions were diluted with complete media to a concentration of 1×10^6^ cells/mL of media.

#### 2.2.2 Print preparation for biocompatibility studies

Disks with a diameter of 15 mm and a height of 1 mm, designed to fit snugly against the base a 24-well plate, were printed in all three representative resins (BioMed, Clear, and BV007a) following the print settings outlined in Table S1. These pieces were divided into “non-treated,” with no further post-treatment, or “treated”. The latter prints were post-treated by soaking in sterile 1x phosphate-buffered saline without calcium and magnesium (PBS, Prod. No. 17-516F, Lonza, USA) for 24 hours at 37°C for BV007a or at 50°C (BioMed and Clear resins) to mitigate cytotoxicity, with similar treatments shown to be effective in previous works.^46^

To compare post-treatment strategies for the BioMed resin, disks were printed as above. The post-treatments included a 24-hr PBS soak at room temperature within a biosafety cabinet; 24-hr incubation in a 37°C cell culture incubator while dry or soaked in PBS; or autoclaving for 30 minutes at 120°C gravity cycle. In all cases, both treated and untreated pieces were rinsed again with IPA, dried, and UV sanitized for an additional 10 minutes before use.

#### 2.2.3 Analysis of cell viability

Aliquots of suspended splenocytes (1 mL, 10^6^ cells/mL) were added to two 24-well plates containing samples of either treated resins or non-treated resins as previously outlined in section 2.2.2. Wells that did not contain any resins were reserved for plate controls. The cell cultures were incubated for 4 hours at 37°C with 5% CO_2_. Following the culture period, the viability of the splenocytes was assessed by flow cytometry using a previously established protocol.^47^ Concisely, 500 μL of the cultured samples were stained using Calcein AM (eBioscience, San Diego, CA, USA) at 67 nM in 1x PBS for 20 min at 37°C. The stained samples were centrifuged at 400 xg for 5 min and resuspended in flow buffer (1 x PBS with 2% FBS), after which 4 μL of 1 mg/mL 7-AAD (AAT Bioquest, Sunnyvale, CA, USA) was added. Calcein-AM single stains were prepared using live cells, and 7-AAD single stains were prepared using cells pre-treated for 20 min with 70% ethanol added in a 1:1 v/v ratio to the culture. Additional controls included unstained cells and an ethanol-treated double-stained control. All samples and controls were run on a Guava 4-color cytometer (6-2L) and analyzed with Guava^®^ InCyte™ Software. Live cells were defined as being high in Calcein-AM and low in 7-AAD signal, while dead cells were defined as the inverse.

#### 2.2.4 Analysis of the viability after direct and indirect contact with treated resin

Indirect contact was defined as cell culture in media that had been conditioned by incubation with printed resin, whereas direct contact was defined as cell incubation in physical contact with the printed resins. To test indirect viability, treated BioMed disks were prepared for cell culture as noted in section 2.2.2. Following treatment, the disks were added to a 24-well plate and incubated in complete media for 24 hours at 37□C. After incubation, 1 mL of suspended splenocytes at 10^6^ cells per mL were spun down and brought back up in 500 μL of resin-conditioned media. All samples were then cultured for 45 minutes, 4 hours, and 24 hours. Viability was analyzed as in section 2.2.3 to determine the percent of live cells present for each sample.

Direct viability was tested in a similar manner. Treated BioMed disks were added to a 24-well plate, and 1 mL aliquots of suspended splenocytes at 10^6^ cells per mL in fresh complete media were added to sample and control wells. Viability was analyzed after 45 minutes, 4 hours, and 24 hours.

### 2.3 Characterization of material properties of printed pieces

#### 2.3.1 Autoclave compatibility and heat tolerance

To test heat stability of printed pieces, small pieces with square channels (as used for print resolution tests, .DWG files are included in the supplement) were 3D printed in each of the three resin types using settings from Table S1 and autoclaved at 120°C for a 30-minute gravity cycle. Following autoclaving, the pieces were visually evaluated for cracks, delamination, or other alterations to the original design. Similar tests were conducted by leaving the prints overnight in ovens at 37°C, 70°C, and 120°C and then visually assessing the prints for discrepancies after 1, 3, and 7 days.

#### 2.3.2 Autofluorescence

Disks with a diameter of 15 mm and a height of 1 mm were printed in each representative resin. A square piece of PDMS was used as a control. All images were collected on a Zeiss AxioZoom macroscope (Carl Zeiss Microscopy, Germany) with Zeiss filter cube channels including Cy5 (Ex 640/30, Em 690/50, Zeiss filter set #64), Rhodamine (Ex 550/25, Em 605/70, #43), EGFP (Ex 470/40, Em 525/50, #38), and DAPI (Ex ~320-385 nm, Em 445/50, #49). A 500 ms exposure time was used for all images. Following imaging, analysis was performed using Image J v1.530 (imagej.nih.gov). On each image, three 1 × 1 in^2^ regions were analyzed for mean gray value in each channel. Background regions were also measured from the borders in each image (outside of the printed parts) and subtracted from each sample measurement individually. The mean gray intensity was calculated for each resin piece and the PDMS control; higher mean gray intensities represented higher autofluorescence of the pieces.

#### 2.3.3 Optical clarity

Disks with a diameter of 15 mm and a height of 5 mm were printed in the FormLabs Clear resin following the print settings listed in Table S1. Several post-processing methods were compared to determine which had the greatest improvement on optical clarity of printed devices. These included printing on glass, a nail polish coating, a resin coating, sanding, and buffing the pieces. A non-processed piece was used as a control, and a glass slide (0.17 mm thick) was used as a benchmark for optimal material clarity.

To prepare the printed-on-glass piece, a print was set up as previously described by Folch, et al.^38^ First, small drops of resin were applied to the baseplate using a transfer pipette. A large cover glass with dimensions of 1.42” × 2.36” and a thickness of 0.13-0.17 mm (Ted Pella, Inc. USA) was attached to the baseplate by lightly pressing the slide over the resin then using a UV flashlight to quickly cure the resin between the slide and the baseplate. In the printer software, the initial layer height (“gap height”) was increased by the thickness of the slide (1.7 mm) prior to printing to account for the change in the z-position of the first layer. After printing, the piece was removed from the glass slide with a razor blade and post-cured typically. The glass slide and adherent resin drops were also easily removed with a razor blade.

For acrylate coating, a baseplate-printed piece was coated with generic clear nail polish from a convenience store. The top was coated using the polish applicator, allowed to dry for ~15 minutes, and the process was repeated on the bottom of the piece. Similarly, a pipette tip was used to apply a thin layer of FormLabs Clear resin to a separate piece on both the top and bottom, with both sides being UV cured for 10 minutes.

For the sanding method, 3M WetorDry Micron Graded Polishing Paper (ZONA, USA) was used. The piece was sanded on both sides starting at a 30 μm grit paper and followed by 15 μm, 9 μm, 3μm, and finally 1 μm grit. Moderate pressure was used to press the piece into the polishing paper in a circular motion to smooth the surface of the piece. Similarly, a generic 4-sided nail buffer (similar products, Walmart, USA) was used to evaluate the impact buffing could have on the printed piece.

Following post-processing, all pieces were imaged to determine optical clarity. All images were collected in the brightfield under transmitted light on a Zeiss AxioZoom macroscope (Carl Zeiss Microscopy). The intensity of light that passed through each piece was measured using Image J for three 1 × 1 inch^2^ sections on each image. The average intensity and standard deviation were recorded for n = 3 regions per sample, with the background subtracted from each measurement individually. The relative transmittance, *T*, of each sample was calculated according to Equation 1,

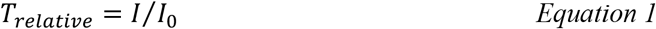

where *I* is defined as the average mean gray intensity of the sample, and *I*_*0*_ is defined as the mean gray intensity of a glass slide. Error was propagated using Equation 2,

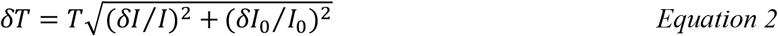

where δ is the standard deviation.

## 3. Results and Discussion

### 3.1 Selecting a resin based on materials properties

Design of a successful 3D printed bioanalytical tool begins with selection of a suitable resin, a process that currently requires compromises. Ideally, the resins used to 3D print bioanalytical microfluidic devices would be compatible with all cell types, be able to produce milli- and microfluidic sized internal features without mechanical defects and meet imaging requirements of having low background fluorescence and high optical clarity when needed. There has yet to be a commercial resin, however, that integrates all of these ideal properties. Custom resin formulations may offer improved performance, at least for laboratories prepared to produce them consistently and tweak them for the intended instrument and application.^31,38,48^ Nevertheless, focusing on enhancing one feature (e.g. cytocompatibility) usually results in compromising on another (e.g. print resolution). Because of this, it is useful to understand the key components of resins before choosing a material to work with.

We have found that many polymeric resins suitable for microfluidics fall into one of three categories based on the use case for which they were designed: biocompatible, optically clear, or high microscale resolution. Though specific resin components differ, resins in the same category often have similar material properties. Therefore, for this work, one test resin from each of these categories was chosen as a case study, and these are listed in Table 1 along with important intrinsic properties. FormLabs BioMed Clear V1 (FormLabs, USA) is a representative “biocompatible” resin with a USP Class VI biocompatibility rating, where it is approved for contact with live mucosal membranes and skin tissue for >30 days.^49^ The FormLabs Clear V4 resin (FormLabs, USA) is representative of a material that offers increased optical clarity. MiiCraft BV007a Clear resin (CADworks3D, Canada) is a low-viscosity resin designed for high print resolution specifically for microfluidic devices. Researchers using one of the dozens of other available resins may use these properties to identify the extent to which it falls into one of these categories and thus predict performance of the resin for the intended use. When protected from ambient light, all resins tested were stable in open vats at room temperature without noticeable variation in volume or viscosity for at least two months.

**Table 1:** Properties of representative SL resins that inform material choice.

| Resin Type | *Biocompatible* | *Optically Clear* | *High Resolution* |
| --- | --- | --- | --- |
| Representative resin | FormLabs BioMed | FormLabs Clear | MiiCraft BV007a |
| Price (USD)* | \$349/L | \$149/L | \$510/kg<sup>†</sup> |
| % Additives in composition* | <2% | <0.9% | 10-15% |
| Viscosity at RT (mPa•s)* | 1350 | 850-900 | 75-100 |
| Heat Stability<sup>‡</sup> | 120°C ~24hr<br>70°C ≥ 7 days<br>Autoclavable | 70°C ≥ 7 days | 37°C ≥ 7 days |

\*As of the date of print, all information is currently available from the vendors and material safety data sheets and are relisted here for convenient comparison.<sup>50-53</sup>
<sup>†</sup> For BV007a, density is 1.04-1.10 g/mL at room temperature, so 1 kg is ~1 L
<sup>‡</sup> Measured in this work

Resins for SL printing are comprised of a photocrosslinkable polymer base, photoinitiators to initiate crosslinking, and (optionally) additives such as photoabsorbers, dyes, and plasticizers.^54–56^ Common photoinitiators such as Irgacure compounds or lithium phenyl-2,4,6-trimethylbenzoylphosphinate (LAP) are activated by UV (365 or 385 nm) or violet (405 nm) light.^55^ While commercial photocrosslinkable resins for SL printing usually keep their exact compositions as a trade secret, the fundamental chemistry can be found in MSDS documentation. All three test resins in this work contained monomers and oligomers of an acrylate or methacrylate polymer base.^50,51,53^ Many SL resins for microfluidics are fairly similar in their basic composition and can be used across different printers, especially with printers that allow for exposure setting adjustments.

Resins may be formulated with additives to achieve a large increase in print resolution or other desired properties. For example, addition of photoabsorbers reduces the effect of scattered light between layers and dramatically improves z-resolution.^32^ Addition of plasticizers lowers the viscosity of the pre-cured resin, which improves print resolution of hollow internal features by facilitating drainage of uncured resin from the feature during printing and cleaning steps. MiiCraft’s microfluidic BV007a resin, with the highest percent of additives, has a manufacturer-reported viscosity about 10-fold lower than that of the other listed resins (Table 1), and superior resolution for microscale channels. In highly viscous materials like the BioMed resin, undrained resin is easily retained inside the internal features, where it may be crosslinked by light that has been transmitted or scattered in excess, especially as subsequent layers of resin are cured above the hollow features.

On the other hand, photoinitiators and plasticizers, as well as acrylate monomers, can increase cytotoxicity with various cell types due to factors including oxidative stress, enzymatic inhibition, and lipophilic reactions with cell membranes.^56–59^ Deliberately leaching these toxic components from the printed materials after post-curing can increase cytocompatibility.^29,30^ However, we have found that in some cases it also decreases material stability, causing cracking or a decrease in material strength and flexibility as the plasticizers are removed. This problem was particularly prevalent with BV007a prints, which peeled apart when leached in PBS for longer than 48 hours, presumably due removal of plasticizers that were essential to the structural stability of the print. For this reason, selecting a more biocompatible material (e.g. with fewer leachable toxic additives) to begin with may better address this problem when working with sensitive cells.^48^

As heat stability is an important factor for chips that will be subject to autoclave sanitation or extended cell culture, we tested the heat stability of each resin (Table 1). The FL BioMed material was autoclavable and also stable overnight at 120°C, allowing for thorough sterilization should any prints need to be reused or prepped for use with live biomaterials. In contrast, MC BV007a withstood mild sanitation procedures, e.g. alcohol rinses or UV sanitation, but high heat (>50°C) delaminated the material over time, although it was stable for 7-day incubation at 37°C when dry, or for 48-hrs in PBS. The FL Clear resin was found to withstand conditions heating conditions up to 70°C for 7 days dry or in solution and also mild sanitation procedures (i.e. UV curing, solvent rinses).

### 3.2 Physiochemical and environmental factors that influence print resolution

In addition to resin composition, the functionality and printability of a piece are also influenced by other factors that affect photopolymerization and thus the resolution and the mechanical integrity of the part (Fig.1).

#### 3.2.1 Choosing wavelength and exposure settings to mitigate light scattering and bleed-through

Internal features such as enclosed microchannels are imperative to many 3D printed microfluidic devices. Most commercial resin materials yield features at the scale of millimeters or hundreds of micrometers on standard printers with 30-40 μm pixel size, and it is possible to improve the print resolution of internal features through strategic selection of resins, light sources and exposure settings, and number of repeat exposures.^32^ The central requirement is to avoid unintentional crosslinking of uncured resin inside the feature, which would lead to blocked features.

Light source wavelength, intensity, and exposure time have a large impact on resolution by modulating on the rate and extent of crosslinking.^31,32,60^ Common light sources include a laser, LED, or UV lamp, and typically emit at 385 nm or 405 nm in commercial printers.^61^ Though most commercial resins can be printed at either wavelength, more efficient reactions are achieved by matching the wavelength of the light source with the excitation and absorbance peaks of the photoinitiators and photoabsorbers, respectively. Doing so decreases the exposure time and intensity required to achieve crosslinking and reduces unwanted scattered and transmitted light through print layers, as documented by Nordin and Wooley^32^ as well as Pontoni.^61^ To briefly demonstrate the impact of wavelength on resolution of internal features, test pieces containing six internal channels of decreasing square cross-section (0.2 - 1.2 mm side-length; Fig. 2A) were printed in FL Clear resin, using either a 405-nm or 385-nm printer (Fig. 2B vs 2C). The X,Y-resolution (effective pixel resolution) was similar between the two printers, at 30 and 40 μm for the 405- and 385-nm printers, respectively. In this resin, the crosslinking reaction was more efficient with the 385 nm light source, enabling reduced light dosage (shorter times and lower intensities; Table S1), which assisted in diminishing bleed-through light allowing uncured resin to drain more easily from channels. Consistent with this, channels were printable at sizes ~0.2 mm smaller with the 385 nm printer (Fig. 2C) versus the 405 nm source (Fig. 2B).

**Fig. 2.**
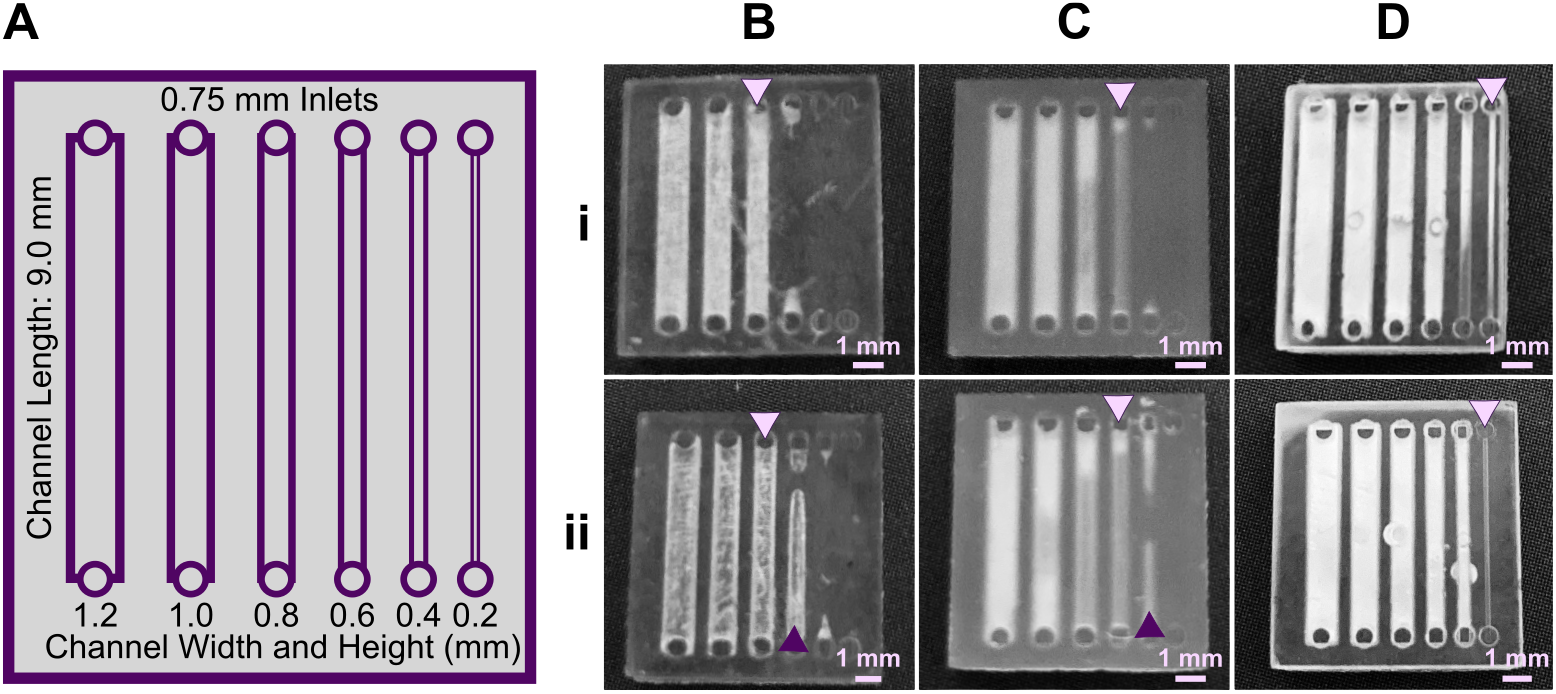
The resolution of internal features can be increased by making changes to the light source, resin viscosity, and print layer height. **(A)** A test piece had six internal channels, 9-mm long with 0.75 mm diameter inlets and varied channel cross-sections as noted. It was printed with **(B-D)** (**i**) 50 μm layer height or **(ii)** 100 μm layer height, as follows: FormLabs Clear resin was printed with a (**B)** 405 nm or (**C**) 385 nm light source; (**D**) for comparison, a low viscosity resin, BV007a, was printed at 385 nm. Other settings were left unchanged, with the exception of increasing exposure times slightly for some 100-μm prints (Table S1). Resolution was determined visually by observing the smallest channel cross-section that could be printed fully (pink arrows) or partially open (purple arrows).

An additional print setting that affects resolution is layer height, which sets the thickness and number of layers that must cured directly above hollow features, known as overhang layers. Each overhang layer, though required to close off the top of the feature, is a chance for light to unintentionally penetrate or scatter into the uncured resin that is trapped in the hollow space, potentially crosslinking it. Increasing the layer height increases layer thickness and reduces the number of overhang layers mitigating some bleed-through curing. This setting can be modified on most printers during the file slicing step when converting a design file into a printable file and works well for designs without strong diagonal features in the z-direction.^63^ Using the FL Clear resin, doubling from 50-μm (Fig. 2i) to 100-μm layer height (Fig. 2ii) improved the print resolution of interior channels in the test piece to partially open the next smaller channel (an improvement of <0.2 mm, Fig. 2ii, purple arrows). Therefore, simply decreasing the number of overhang layers decreased the degree of overexposure or bleed-through light and improved print resolution, though not as much as changing the light source.

#### 3.2.2 Lower viscosity improves drainage from internal channels

As noted in Section 3.1, the need to drain uncured resin out of hollow features during both printing and cleaning means that resin viscosity has a major impact on the resolution of internal features. To demonstrate this, we compared the resolution of FL Clear resin (Fig. 2C) to MC BV007a (Fig. 2D), which have viscosities of ~900 mPa s and ~100 mPa s, respectively (Table 1), using the 385 nm light source. The FL Clear resin retained uncured resin in the channels during printing (visible when the device was removed from the printer), and produced open channels only down to 0.6 mm under these conditions. In contrast, no residual BV007a resin was observed prior to rinsing the channels, and the print yielded a resolution of 0.2 mm, the smallest size tested, thus confirming the significant benefit of low viscosity for channel resolution.

In summary, with its ability to print efficiently at 385 nm and to drain easily with low viscosity, BV007a provided the best print resolution, with 0.2-mm channels printing cleanly at a 50-μm layer height. Printing with the Clear resin, however, achieved nearly 0.4 mm channels if used with a 385 nm light source and 100-μm layer height. While the specifics of printability will change for each design, we found that the light source, material viscosity, and layer height each provide opportunities to increase print resolution of interior microchannels.

#### 3.2.3 Influences of print environment on photopolymerization

Finally, the humidity and temperature within or surrounding the printer can also impact how well the photopolymerization reactions take place. Many resins are recommended to be printed at relatively low humidity (20-40%), as photopolymers are often hygroscopic.^64^ We have observed that when the humidity increases (averaging 45% in the summer in our laboratory, sometimes higher than 50-60%; versus ~30-35% in winter), print failures became more common: print layers delaminated or base layers did not remain attached to the baseplate. Keeping printers and resins in an environment with regulated humidity should be considered, e.g. dehumidification. We also found that increasing the power of the light source by 5-10% partially compensated for humidity increases.

Ambient temperature can also influence how well a material prints as it directly impacts the viscosity of the material and reaction times. This topic was explored in depth by Steyrer, et al. in an evaluation of hot vs. room temperature lithography.^65^ In general, many resins tend to produce better print resolution in a warmer environment (above 20°C) due to lower viscosity and more efficient polymerization.^65,66^ For this reason, some printers allow for control over vat temperature. For printers without such control, we have found that decreases in ambient temperature (e.g. from 23°C to 20°C) can be compensated by increasing exposure time, usually by a few seconds depending on the resin and other factors, to overcome the decreased reaction efficiency. Care should be taken, though, as increasing exposure times may lead to overcuring.^65^

### 3.3 Print and design considerations for common microfluidic features

In this section, we discuss design principles for troubleshooting common print failures and improving the resolution of 3D printed microfluidic channels, along with the role of light exposure and printer settings in controlling print quality.

#### 3.3.1 Selecting a print orientation

The build orientation and design of a part impact how the print experiences gravitational strain and other mechanical stress during printing. Print orientation techniques are a common topic for 3D printing blogs and education resources.^67–69^ In general, considerations for choosing a suitable build orientation include baseplate adhesion, location of part functionalities, the use of build supports, and fabrication time. Baseplate adhesion is usually strongest when the largest surface area of a design can be placed flush against the baseplate (Fig. 3A–D, i), which increases the overall stability of piece during the printing process. A slightly rough surface allows prints to adhere better to the plate while printing with lower base layer curing times, although too much adhesion makes removal difficult and can lead to breakages. Location of part features also influence how to orient a print. In this example, orienting the reservoir sideways (ii, iv) caused distortion of its walls or damage from supports, and inhibited resin drainage from the channel. Surface roughness or clarity were also influenced by the direction in which the layers were built: clarity and smoothness are maximized by orienting so that the face of the chip is printed in one layer (i, iii), and reduced by gravitational strain (Fig. 3C, iii) or too many print supports (Fig. 3D, iii). The latter can also contribute to overexposure and light scattering, causing features to fill in (Fig. 3D, ii-iv). Finally, as each additional layer adds cure time to the print, and supports add post-processing time, it is useful to minimize the use of supports while printing if possible, and to select print orientations with fewer layers to create a more efficient printing process.

**Fig. 3.**
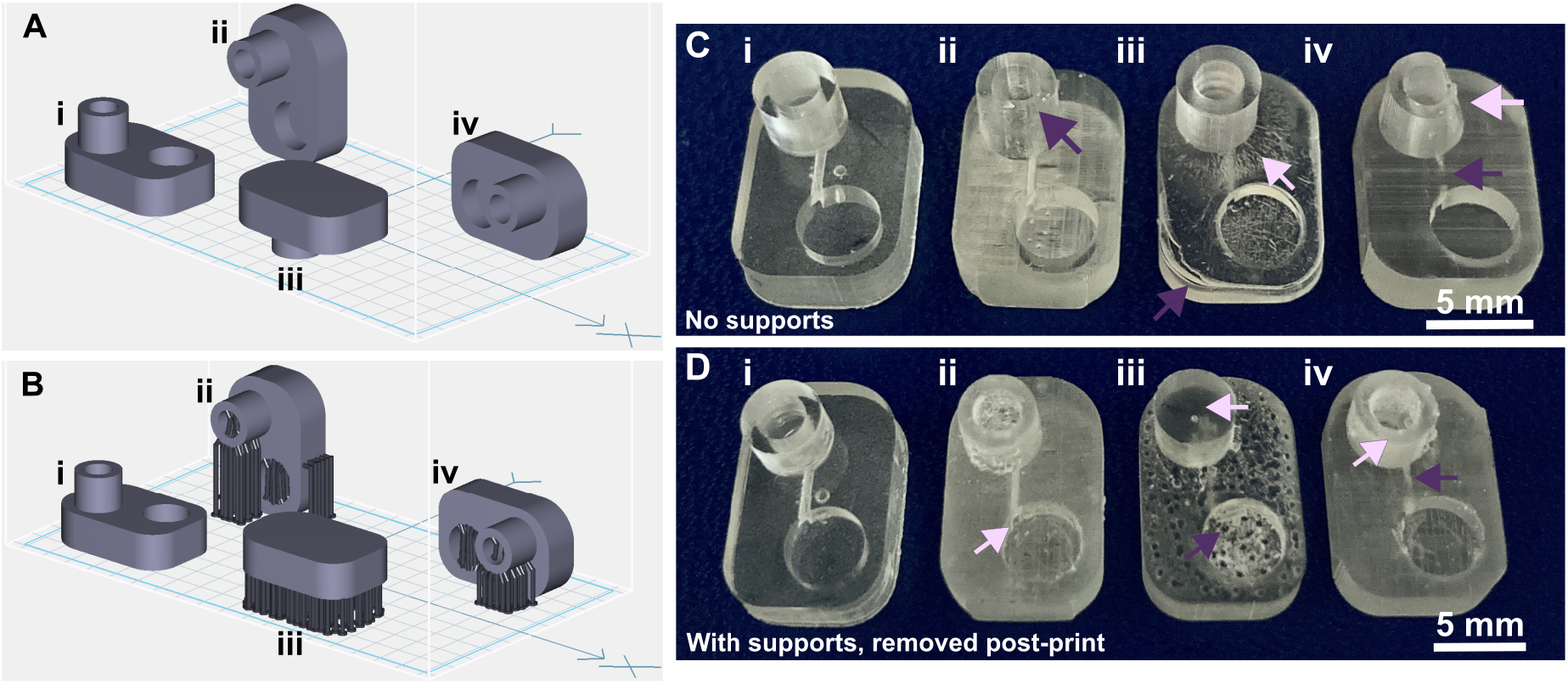
Orientation of prints on a build plate impacts the print resolution and optical clarity. A microfluidic design containing a reservoir, channel, and chamber is shown in the MiiCraft Utility slicing software in four orientations (**A**) without supports and (**B**) with supports: (i) base-down, (ii) vertical, (iii) top-down, and (iv) horizontal. Blue grid represents the base plate. Supports were added to all areas using default settings in the slicing software. (**C, D)** Photos of pieces printed in BV007a resin with the MiiCraft Ultra 50 printer, (**C**) printed without supports, and (**D**) printed with supports, with 2 minutes of support removal. Surface roughness and clarity variation is evident between the 4 orientations. Defects were noted as follows: In **C**: (i) no defects, (ii) overhang drooping, (iii) delamination (purple) and stress fracturing (pink), (iv) blocked channel (purple), overhang drooping (pink); in **D**: (i) no defects, (ii) stress fracturing, (iii) damage from supports (purple) and filled chamber (pink), (iv) filled channel (purple) and support damage (pink).

#### 3.3.2 Reducing mechanical strain in wells, cups, or ports

In microfluidic design, particularly for bioassays, wells or cup-like features are often used as reservoirs or ports. If not properly designed, these wells often fall victim to print failure.^70–72^ Like other photocrosslinkable polymers, many SL resins shrink 1-3% by volume upon crosslinking which may induce mechanical strain.^55,71,73^ If the walls of a print are thinner at some points and thicker at others, the thicker regions experience greater shrinkage and may generate defects.^71^ Weak structural points may form when thin walls, sharp corners, and sharp edges (~90° angles) are used in a design, and these may not hold up well in the printing process.^74,75^ Wells or cup-like features, for example, may experience “cupping,” which occurs when hollow features become damaged during printing due to the formation of a pressure differential (Fig. 4). This effect is due to a low pressure region, or suction, formed within the feature as the print is peeled or pulled away from the Teflon sheet after each layer is exposed, leading to cracks and/or holes at weaker structural points in the design as it caves inward under the surrounding pressure (Fig. 4B).^70^ Such defects may cause leaking when the printed well is later filled with fluid.

**Fig. 4.**
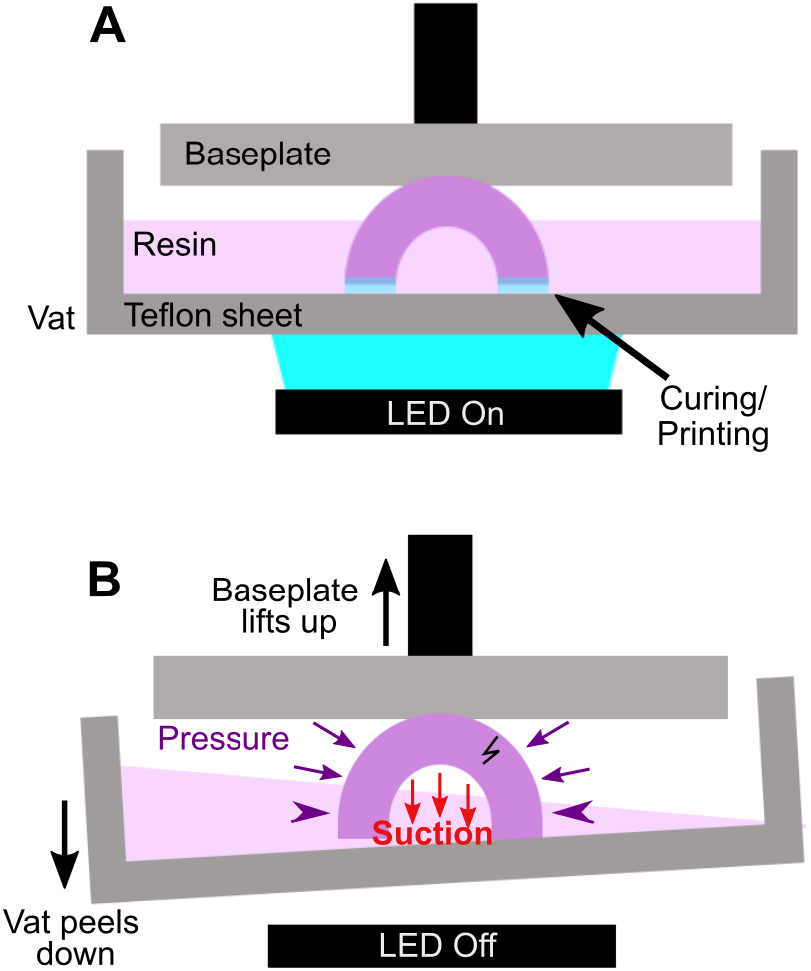
Schematic of “cupping” damage during the printing of a hollow, cup-like feature. (**A**) During printing, a UV light source (LED) forms a new crosslinked layer of resin flush against the Teflon sheet. This design is an inverted bowl shape; supports are not shown for clarity. (**B**) After a layer is finished printing, the print is peeled away from the Teflon sheet, e.g. by pulling the vat down and/or the baseplate up. This process creates a region of suction within the hollow cup feature; the surrounding pressure, now higher than the pressure within piece, may form a stress fracture on the print.

As a demonstration, a hollow well printed in a square base with 90° square angles and thin walls broke routinely under the pressure build up from cupping (Fig. 5A). We tested the impact of various strategies to reduce mechanical strain in the design, drawing upon engineering principles.^74,75^ The square exterior, with its varied wall thickness around the radially symmetric well, contributes to an uneven stress distribution during resin shrinkage. Increasing the thickness of the base and wall and filleting the connecting edge at the base of the well reduced the risk of cracking the base of the print, but not the appearance of small holes at the base of the well (Fig. 5B). Making the thickness of the walls more uniform around the well-like feature, by rounding the exterior corners of the feature either partially (Fig. 5C) or fully (Fig. 5D) reduced the strain unequal shrinkage as expected, with full rounding producing a well feature with no holes or other leaks.

**Fig. 5.**
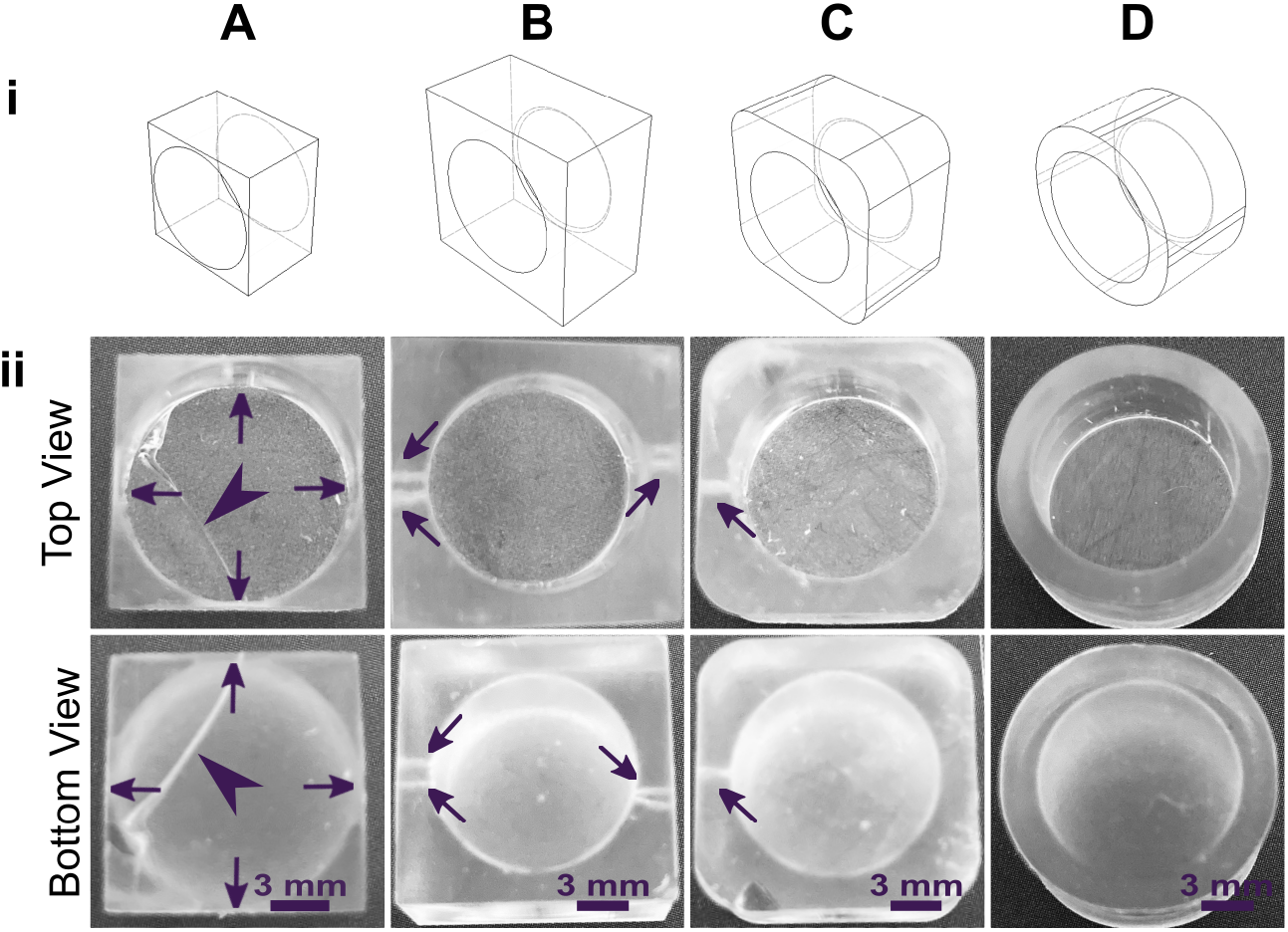
Strengthening the design around well-like features decreases the impacts of cupping and resin shrinkage. (i) Illustrated computer-aid designs, and (ii) photos of the top and bottom views of the corresponding print. All pieces were printed in FormLabs Clear resin. Well **A** had thin walls, thin base, and strain from 90□ connections at the bases and the sides. Well **B** had thicker surrounding walls and a thicker, filleted base, but retained 90□ outer corners. Wells **C** and **D** further reduced the strain by rounding out the external edges. All pieces were qualitatively evaluated, with the absence of cracks (arrowheads) and pinholes (shown by arrows) indicating a strong design.

#### 3.3.3 Improving print resolution of interior channels via device design

Similar to the effect of layer height during printing, we predicted that changing the device design itself to reduce the number of UV exposures over the channel or otherwise facilitate drainage of uncured resin would improve resolution. To demonstrate, we again selected the FL Clear resin due to its high viscosity and transparency, properties that result in frequent bleed-through curing. The number of overhanging layers was controlled by repositioning the channels in the z-direction (Fig. 6), using a fixed layer height of 50 μm. Square channels that were printed near the base of the print, with 1.5 mm thick overhangs and thus 30 overhang layers, printed cleanly only down to 1.0 mm width (Fig. 6B). Decreasing the overhang thickness to 0.5 mm (10 overhang layers) improved the resolution by ~0.2 mm (Fig. 6C), consistent with the prediction. Additionally, adding drainage holes further improved print resolution of long channels that otherwise did not drain (Figure S1).

**Fig.6.**
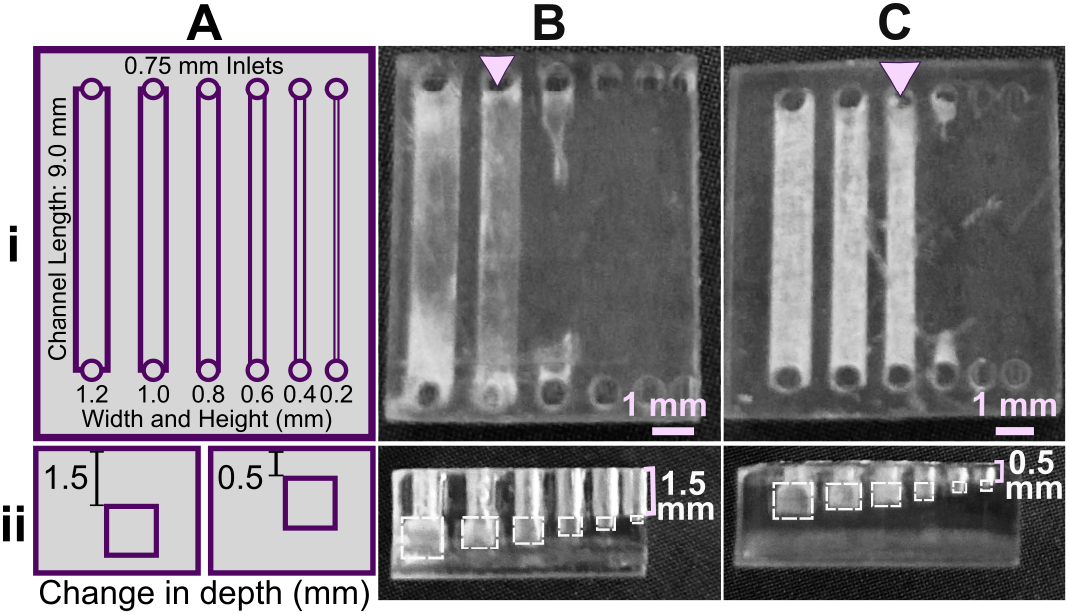
Reducing the number of overhang layers in the chip design improved the resolution of internal features. (**A**) Schematic of the test piece with six internal channels, in (**i**) top view and (**ii**) side view. (**B**,**C**) Channels were printed with (**B**) 1.5 mm or (**C**) 0.5 mm overhang thickness, and imaged from (**i**) top and (**ii**) side. The square channel is traced with dashed outlines; the features visible above the channel in the side view are inlets and outlets. All pieces were printed in FL Clear resin with a 405 nm light source (settings in Table S1). Resolution was determined visually by observing the smallest channel cross-section that could be printed fully (pink arrows).

### 3.4 Cytocompatibility of resins with primary cells

Within this section, we discuss actionable steps towards improving resin cytocompatibility, as it is typically the most challenging issue for 3D printing microfluidics to be used with live cultures and tissues. Many recent publications have addressed this issue, with most solutions focused on preventing potentially toxic resin components from reacting with cell and tissue cultures. Methods include coating the prints to reduce direct contact with cells;^24^ minimizing unbound toxins via autoclaving, overcuring, or pre-leaching toxic substances;^29^ or producing resins with more biocompatible photoabsorbers.^38,76,77^ Leaching in particular is convenient and consistent, especially for treating internal print features that may not be directly accessible to over-curing. Therefore, here we first compared biocompatibility across the representative commercial resins, with and without a standardized, leaching-based post-treatment in heated saline (see Methods; Fig. 7A).^18,29^ Primary, naïve murine splenocytes were chosen for the viability tests as they are typically more sensitive to small changes in their culture environment than hardier, immortalized cell lines.^78,79^

**Fig. 7.**
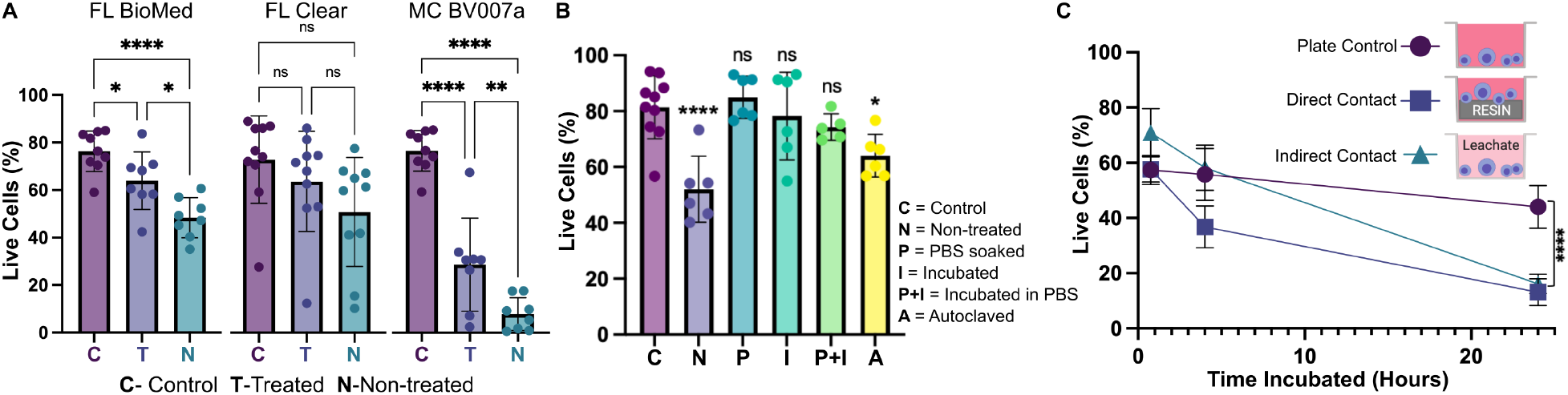
Viability of primary murine splenocytes in contact with 3D printed materials. Primary splenocytes from male and female mice (N_mice_ = 2 per experiment) were evaluated by flow cytometry after live/dead staining with Calcein-AM/7AAD. (**A**) Cell viability after 4 hr of direct physical contact with PBS+incubation treated (*T*) or non-treated (*N*) resins compared with well plate control (*C*). Treated BioMed and Clear prints maintained relatively high viability compared to the well plate, while BV007 did not. Bars show mean ± SD. One-way ANOVA per data set, *ns* > 0.06, \**p* > 0.01, \*\**p* = 0.01, \*\*\*\**p* < 0.0001. (**B**) Multiple post-treatment methods (PBS, incubation at 37 °C, PBS+incubation, and autoclavation) were evaluated using BioMed resin and culturing for 4 hours with direct contact. Bars show mean ± SD. One-way ANOVA, *ns* > 0.2, \**p* > 0.02. \*\*\*\**p* < 0.0001. (**C**) Viability of direct contact (i.e. culture with resin) and indirect contact (i.e. culture with resin leachate) of cells with treated BioMed prints at 45 min, 4 hours, and 24 hours showed a decrease in viability over time. Values show mean ± SD. One-way ANOVA at final time point, \*\*\*\**p* < 0.0001.

We found that for experiments under 4 hours, overnight heat and saline leaching treatment was sufficient to increase viability of primary murine splenocytes in contact with 3D printed materials (Fig. 7A). Splenocytes cultured in direct physical contact with either of the treated FL resins retained viability high enough (>60%) after 4 hr to enable short on-chip experiments, whereas untreated materials resulted in lower viability on average. For the MC BV007a resin, the treatment did improve viability over the non-treated resin, but the resin was still largely cytotoxic for these cells at 4 hours. The higher cytotoxicity is consistent with the greater quantity of potentially toxic additives in BV007a compared to other resins (10-15%, Table 1). Furthermore, BV007a had lower heat stability and could not be leached at the same temperature as the FL resins without mechanical damage (Section 3.1). Therefore, for short experiments with primary cells in suspension, we concluded that the Biomed or Clear resins are more suitable than BV007a and should be pre-treated to minimize release of toxic components into the culture.

Next, we compared a number of common leaching treatments head-to-head: heat (37°C incubation, 24 hr), saline (sterile PBS soak at room temperature, 24 hr), a combination of the two (sterile PBS soak at 37°C, 24 hr), or autoclaving (gravity, 120°C, 30 min) to leach unbound toxins out of solid prints (Fig. 7B). Using the BioMed resin, we found all overnight saline (PBS) and incubation (37°C heat) treatments increased splenocyte viability at 4 hours, with non-significant differences between the live plate control and treated materials. Autoclaving was slightly less effective, and though this treatment takes less time (~45 minutes), it is not recommended, at least for this resin.

Finally, as many experiments require longer term culture than just 4 hr, we tested splenocyte viability over time when cultured in direct physical contact with treated FL BioMed resin or in “indirect contact,” i.e. cultured in contact with media that was pre-conditioned by incubation with the resin. Direct contact with resin is likely to occur in a fully integrated bioanalytical microchip with on-chip cell culture, while indirect contact may occur when microdevices are used to prepare media or drug solutions (e.g. mixers or droplet microfluidics) or to deliver media components to cultures (e.g. flow channels or bubble traps). We again found that splenocyte viability was high for both contact conditions within 0-4 hr, but decreased overnight (24 hrs) for both contact conditions compared to plate controls (Fig. 7B). Others have reported longer culture times for hardier cell lines.^18,25,27–29,33,38,80,81^ This result indicates that at least for primary murine splenocytes and perhaps for other fragile cells, more work is needed to identify best practices for post-print treatments and/or more biocompatible resin formulations.

### 3.5 Optical components of resins

Microfluidics is frequently integrated with on-chip imaging. Autofluorescence is a potential limitation of polymeric chips, whereas PDMS is often praised for its optical compatibility. Therefore, we quantified the autofluorescence of each of the representative resins in four standard fluorescence channels. In the Cy5 and Rhodamine channels, all three resins showed relatively low autofluorescence and were comparable to PDMS (Fig. 8). However, UV excitation (DAPI channel) elicited high levels of autofluorescence from the two FormLabs resins. The Clear resin also showed moderate autofluorescence in the GFP channel. Autofluorescence at short wavelengths is consistent with the use of photoabsorbers intended to absorb at short wavelengths.^76^ On the other hand, MC BV007a had negligible autofluorescence in all channels, similar to PDMS. Thus, optical compatibility in the intended fluorescence channels should inform material choice for microfluidics devices.

**Fig. 8.**
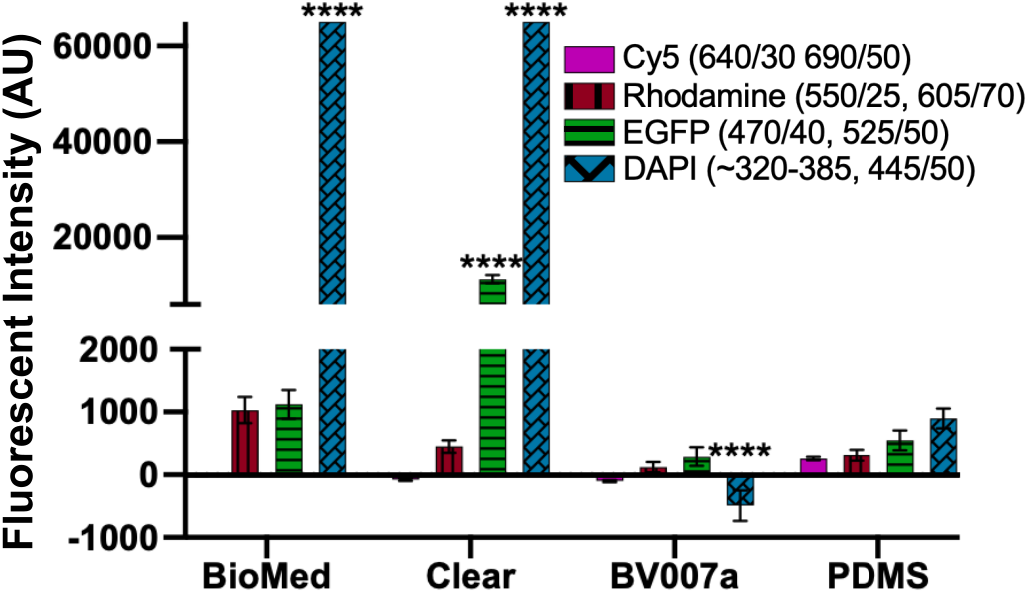
Autofluorescence depended on resin composition and fluorescence filter set. BioMed, Clear, and BV007a prints were evaluated in comparison to PDMS for autofluorescence in Cy5, Rhodamine, EGFP, and DAPI fluorescent channels. Background subtracted mean grey values were analyzed with ImageJ and used to determine the fluorescent intensity of each material. Saturation values were at 65,000 AU. Bars show mean ± SD, N=3 intensity measurements per print. Two-way ANOVA, comparisons shown between resin results versus PDMS for each respective fluorescent channel, \*\*\*\**p*<0.0001.

Optical clarity (transparency) is also important for microscopy and imaging. Various approaches for improving this property have been suggested in published papers as well as on vendor and hobbyist websites (r\3Dprinting, All3DP, etc.).^37–39^ Here we compared 5 of these methods head to head, using the FL Clear resin as a base, to determine which would be best for increasing the transparency of a printed piece (Fig. 9). All methods were compared to a glass slide as a positive control. Pieces that were printed flush against the rough base plate (i.e. without the use of supports or rafts) came off the base plate slightly cloudy in appearance (Fig. 9VI). This cloudiness was intensified when attempting to buff both the base end and the vat end of the print with a typical nail file (Fig. 9VII). Since the nail file did not have a fine sanding grain, it ended up doing more harm than good by producing new scratches on the surface of the print.

**Fig.9.**
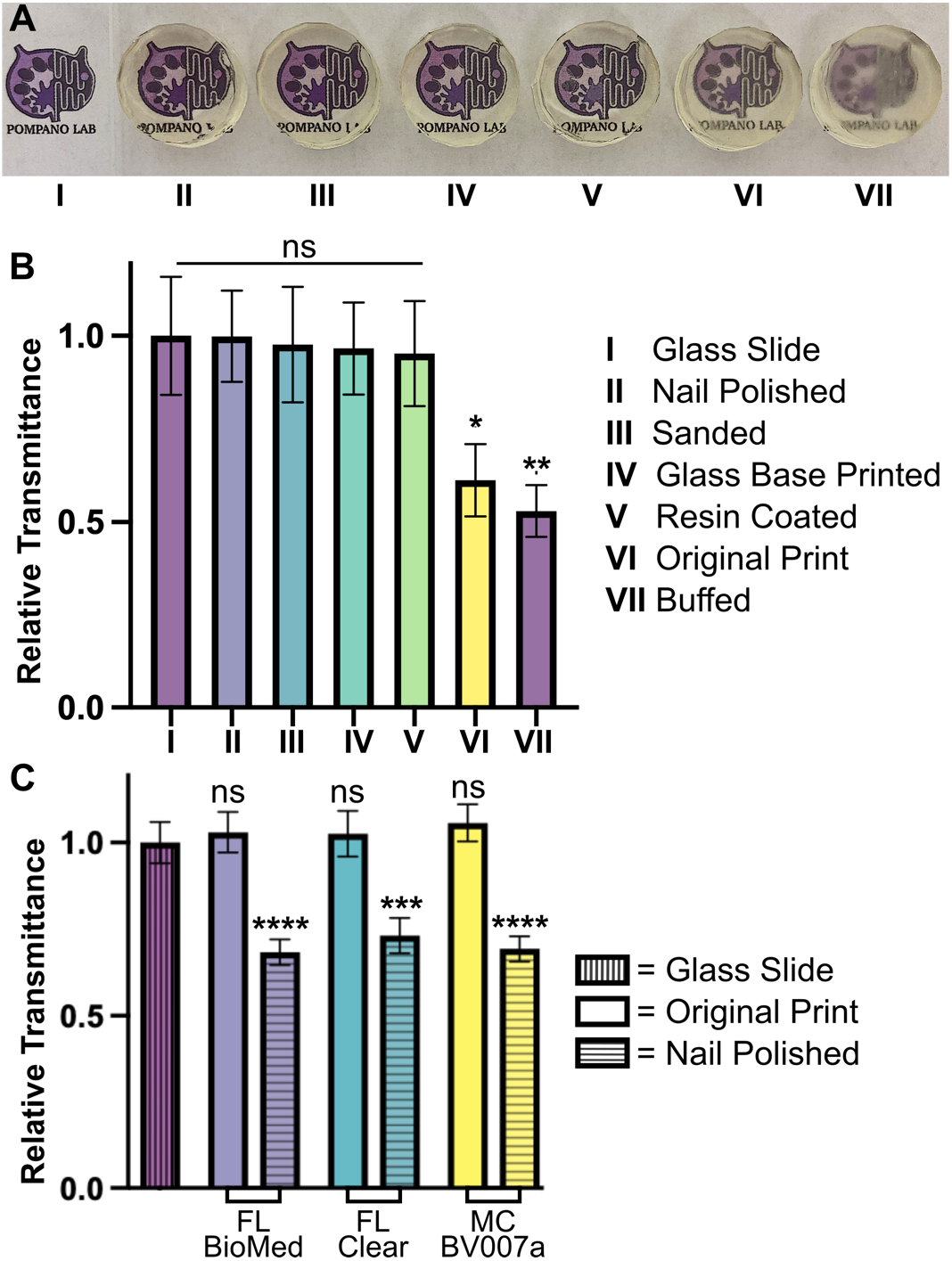
Optical clarity of clear resins was enhanced with post-treatments that achieved material transparency similar to glass. Round prints of FormLabs Clear resin were printed with the MiiCraft Ultra 50 405 printer and post-treated as listed in the legend, labels II -VII. (**A)** The test pieces were positioned over the Pompano Lab logo for qualitative, visual comparison (captured with phone camera). (**B)** Transmittance was evaluated using an upright microscope. Relative transmittance was compared to the average light transmittance through a 0.17 mm thick glass slide (I). Bars show mean ± standard deviation, n = 3. One-way ANOVA, Tukey’s post-hoc test; \**p* = 0.0284, \*\**p* = 0.0067, and ns > 0.05. (**C**) To show reproducibility for different resins, technique II was applied to similar round prints in FL BioMed, FL Clear, and MC BV007a resins and evaluated similarly to panel B. All samples were statistically similar compared to the glass slide control. Two-way ANOVA, Tukey’s multiple comparison test; \*\*\**p*=0.0003, \*\*\*\**p*<0.0001, and ns>0.8.

Several methods proved successful in improving optical clarity. The quickest method that produced glass-like clarity was the nail polish coating of both the base and vat sides of the print (Fig. 9II). The nail polish only took a few seconds to apply and approximately 10 minutes to dry on each side, which introduced a challenge to keep dust out of the polish while drying on an open countertop. Sanding the piece with micron grain sandpaper avoided the issue of dust and the pieces emerged smooth (Fig. 9III) instead of scratched like the nail-filed (“buffed”) piece (Fig. 9VII). Sanding the printed part on at least the baseplate-side is especially recommended if it will not disrupt any print features, as it more easily maintained a uniform surface than the other methods. Printing directly on glass^38^ (Fig. 9IV) had similar impacts to sanding and is recommended for smaller prints that would be easier to remove from a glass slide. This approach requires some caution, though, as attaching glass to a baseplate can cause increased wear and tear to a printing vat. It also requires resetting the initial layer height so that the printer does not lower the baseplate too far into the vat. Resin coating (Fig. 9V) was also helpful for increasing transparency, but it was difficult to achieve without overcuring the rest of the print or under-curing the additional coat, which was also prone to capturing dust. Relative to the glass slide control, most of the post-treated 3D prints were determined to have acceptable transparency. The nail polish technique (Fig. 9II) was tested on all three representative resins and consistently improved the relative transparency of each resin (Fig. 9C). We expect that each of methods I-V will provide improved optical clarity to most transparent or semi-transparent resins.

## 4. Conclusion

In using 3D printing for production of microfluidic devices, compromises and strategic design choices are often required to best match the material and design to the required experiment. After identifying priorities based on the planned experiments, a resin should be chosen that best fits the requirements of print resolution, mechanical stability, cytocompatibility, and optical compatibility, informed by a foundational understanding of material components. If needed, printer settings and device designs can be modified to increase the integrity of printed parts and resolution of interior channels, and post-treatment methods can be used to increase the cytocompatibility and optical clarity of a printed piece. Print stability can be improved by reducing mechanical stress in the design of a piece, and internal feature resolution can be increased by ensuring adequate resin drainage and minimizing the photoexposure of trapped resin, e.g. by reducing the number of overhanging layers. Viability can be improved upon by leaching toxins out of prints prior to application with cells, though there is still a need for more biocompatible options, especially for sensitive primary cells. Optical clarity of parts printed with clear resins can be improved via polishing methods to achieve glass-like transparency. In the future, resins that are high-resolution, cytocompatible, and optically clear will certainly be an area of continued commercial development, and promising PEG-DA based resin formulas have been reported that can be made in the laboratory.^38,76,77^ Meanwhile, the considerations and best practices recorded here can help researchers begin to integrate SL 3D printing fabrication with commercially available products into their microfluidics research. We envision that this guide and its head-to-head comparison of conditions will help streamline the fabrication workflow for researchers who are new to 3D printing within the biomicrofluidic community.

## Supporting information

Autodesk Files for 3D Prints

## Author contributions statement

**H.B. Musgrove:** Conceptualization, methodology, investigation, data curation, formal analysis, visualization, writing – original draft, review and editing **M.A. Catterton:** Conceptualization guidance, methodology, writing – review and editing **R.R. Pompano**: Funding acquisition, supervision, writing – review, editing, and finalization

## Acknowledgements

Research reported in this publication was supported by the National Institute of Allergy and Infectious Diseases under Award No. R01AI131723 and from the National Institute of Biomedical Imaging and Bioengineering under Award No. R03EB028043 through the National Institute of Health (NIH). The content is solely the responsibility of the authors and does not necessarily represent the official views of the National Institutes of Health. The authors would like to thank Amirus Saleheen for his suggestions for post-treating resins in preparation for cell culture, Scott Karas for his assistance with sanding prints, Alexander Ball for his expertise and assistance with flow cytometry experiments, and Michael Ly for his correspondence regarding MiiCraft and CadWorks3D resins and printers.

