## Supplemental Information for "Applied tutorial for the design and fabrication of biomicrofluidic devices by resin 3D printing"

Electronic Supplementary Information for

**Contents:**

**I.** Supplemental Table S1 showing print settings and post-treatments

**II.** CAD files for well and channel designs (Separate files)

**III.** Supporting Figure S1 and accompanying text

### I. Supplemental Tables

**Table S1**: Print and post-cure settings for each resin

Printer Settings for CADworks3D M50 405 nm

| | *Layer Height (μm)* | *Cure Time (s)* | *Peel Speed* | *Gap Adjustment (mm)* | *Base Layers* | *Base Cure Time (s)* | *Buffer Layers* | *Power %* |
|---|---|---|---|---|---|---|---|---|
| *FormLabs BioMed* | 50, 100 | 5.00 | Slow | 0.10 | 1 | 8.00 | 5 | 75 |
| *FormLabs Clear* | 50 | 5.00 | Slow | 0.10 | 1 | 6.00 | 1 | 80 |
| | 100 | 9.00 | Slow | 0.10 | 1 | 12.00 | 1 | 75 |
| *MiiCraft Bv007a Clear* | 50, 100 | 1.15 | Slow | 0.10 | 1 | 9.00 | 1 | 75 |

Printer Settings for CADworks 3D Printer P110Y 385 nm

| | *Layer Height (μm)* | *Cure Time (s)* | *Peel Speed* | *Gap Adjustment (mm)* | *Base Layers* | *Base Cure Time (s)* | *Buffer Layers* | *Power %* |
|---|---|---|---|---|---|---|---|---|
| *FormLabs BioMed* | 100 | 3.00 | Slow | 0.10 | 1 | 8.00 | 5 | 75 |
| *FormLabs Clear* | 50 | 3.50 | Slow | 0.10 | 1 | 7.00 | 4 | 58 |
| | 100 | 4.00 | Slow | 0.10 | 1 | 8.00 | 4 | 60 |
| *MiiCraft Bv007a Clear* | 100 | 1.50 | Slow | 0.10 | 1 | 25.00 | 2 | 100 |

Post-Print Settings

| | *Alcohol Rinse Time (FormWash)* | *Drying* | *Post-cure Time* | *Post-cure UV Power* | *Post-cure Temperature* |
|---|---|---|---|---|---|
| *FormLabs BioMed* | 20 min | Compressed air used for 2 minutes minimum, then let sit until completely dry | 60 min | 39 W 405 nm LED | 50°C |
| *FormLabs Clear* | 10-15 min | | 20 min | | 45°C |
| *MiiCraft Bv007a Clear* | 3-5 min | | 1 min | | RT |

### II. CAD files for well and channel designs

These files (Autocad dwg. extension) contain the designs for the rounded well and resolution test designs printed for Fig. 3-6. They are provided separately.

### III. Supporting Figures

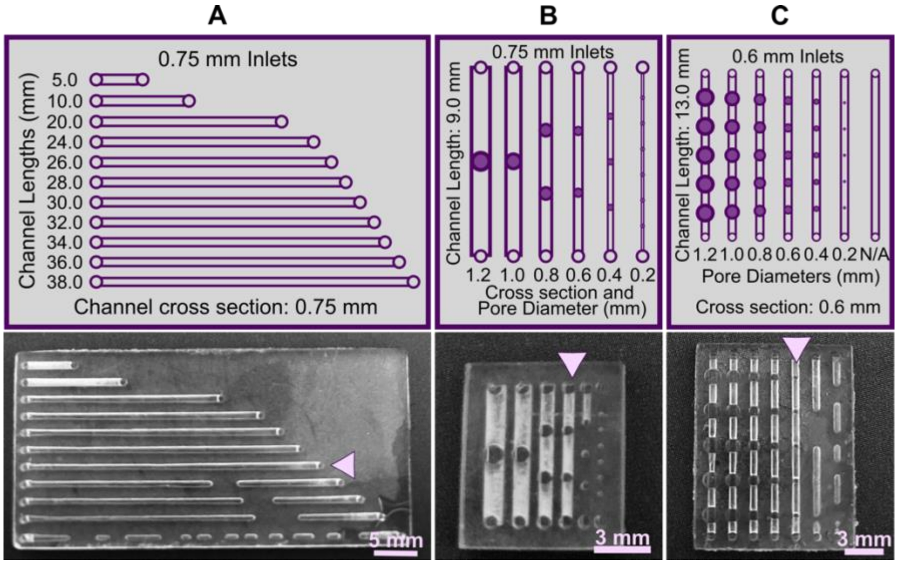


**Fig. S1 Improving drainage and print resolution of microfluidic channels by adding apertures or decreasing channel length.** A series of prints were generated using a printer with a 405 nm light source, 100 µm layer height, and FormLabs Clear resin. Each panel shows a schematic (*top*) and photo of the printed part (*bottom*). (**A**) Channels of varied length from 5.0 – 38.0 mm, with fixed 0.75 mm cross sectional dimension. (**B**) An adaptation of the original test piece with draining pores (filled purple circles) that were of the same width as each channel, ranging from 0.2 – 1.2 mm cross section. (**C**) An adaptation of the original test piece draining pores of varied pore diameter (0.2 – 1.2 mm), over a fixed channel cross section of 0.6 mm. Resolution was determined visually by observing the smallest channel cross-section that could be printed fully in each design (pink arrows).

**Accompanying text:** The ability to print open internal microchannels is limited by the ability for uncured resin to drain from the channel during and after printing. One strategy to improve resin drainage from interior channels is by using shorter channels or adding drainage pores, when feasible in the design. Here, we demonstrated this point and characterized the best conditions for the drainage pores in a series of test prints in the highly viscous Formlabs (FL) Clear resin.  Especially with smaller cross sections (≤0.5 mm), channels were increasingly difficult to drain as the length of the internal channel increased (**Fig. S2A**). Should this kind of blockage occur in a feature, draining pores potentially could be added. Indeed, in the viscous FL Clear resin, pores added to the channels in the test piece improved print resolution from 1 mm (Fig. 5B) to 0.6 mm (**Fig. S2B**); in this experiment, we arbitrarily added more pores to channels with smaller cross sections to ensure sufficient drainage. We found that it was best to use pores with a diameter the same or slightly smaller than the channel; smaller pores did not provide sufficient drainage and were filled with uncured resin (**Fig. S2C**). A potential concern with this method is that large open pores may cause leakage when the channel is filled with fluid. We have found that in low-pressure flow applications, surface-tension based pinning is sufficient to prevent leakage (data not shown). Should one wish to fill the holes, however, several options exist. We found that pinholes can be filled with uncured resin through capillary action and cured with UV light (data not shown), though care should be taken to not block the internal channels.
